# Heterogeneity of age-related neural hyperactivity along the CA3 transverse axis

**DOI:** 10.1101/2020.08.31.275156

**Authors:** Heekyung Lee, Zitong Wang, Scott Zeger, Michela Gallagher, James J. Knierim

## Abstract

Age-related memory deficits are correlated with neural hyperactivity in the CA1 and CA3 regions of the hippocampus. Abnormal CA3 hyperactivity in aged rats has been proposed to contribute to an imbalance between the normal tradeoff between pattern separation and pattern completion, resulting in overly rigid representations. Recent evidence of functional heterogeneity along the CA3 transverse axis suggests that proximal CA3 supports pattern separation while distal CA3 supports pattern completion. It is not known whether age-related CA3 hyperactivity is uniformly represented along the CA3 transverse axis. We examined the firing rates of CA3 neurons from male young and aged Long-Evans rats along the CA3 transverse axis. Consistent with prior studies, young CA3 cells showed an increasing gradient in mean firing rate from proximal to distal CA3. However, aged CA3 cells showed an opposite trend, with a decreasing gradient from proximal to distal CA3. Thus, CA3 cells in aged rats were hyperactive in proximal CA3, but possibly hypoactive in distal CA3, compared to young rats. We suggest that, in combination with altered inputs from the entorhinal cortex and dentate gyrus, the proximal CA3 region of aged rats may switch from its normal function that reflects the pattern separation output of the DG and instead performs a computation that reflects an abnormal bias toward pattern completion. In parallel, distal CA3 of aged rats may create weaker attractor basins that promote bistable representations under certain conditions.

## Introduction

Elevated excitability in the hippocampus contributes to impaired memory function in aging (Gray and Barnes, 2019; Haberman et al., 2017a; Samson and Barnes, 2013; Yassa and Stark, 2011). Specifically, heightened activation localized to the DG/CA3 regions of the hippocampus is correlated with memory deficits in aging and in prodromal Alzheimer’s disease (AD) such as in patients with amnestic mild cognitive impairment (aMCI) (Bakker et al., 2015, 2012; Reagh et al., 2018; Yassa et al., 2011, 2010). In agreement with those data, augmented neural activity occurs in the CA3 region in aged rats (Haberman et al., 2017b; Maurer et al., 2017; Robitsek et al., 2015; Wilson et al., 2005) and in aged primates (Thomé et al., 2016). Treatments to reduce hyperactivity with low dose administration of the atypical antiepileptic levetiracetam reduced firing rates and IEG markers of hyperactive CA3 neurons (Haberman et al., 2017b; Robitsek et al., 2015) and improved performance on memory tasks in aged rats (Haberman et al., 2017a; Koh et al., 2010). Therapeutic efficacy with levetiracetam treatment has also been demonstrated in aMCI patients, who showed reductions in task-related DG/CA3 hippocampal activation and improved memory performance (Bakker et al., 2015, 2012).

Hippocampal place cells fire in specific locations when an animal explores an environment (O’Keefe, 1976; O’Keefe and Dostrovsky, 1971). The characteristics of CA3 place fields have shown age-related changes. CA3 place cells in aged rats have been described as “rigid’, in that they maintain the same spatial representation abnormally across familiar and novel environments (Robitsek et al., 2015; Wilson et al., 2005). Abnormal DG/CA3 activity in old age has also been linked to a reduced ability to discriminate between similar objects in both humans (Bakker et al., 2015, 2012; Berron et al., 2019; Reagh et al., 2018; Yassa et al., 2011, 2010) and in aged rats (Johnson et al., 2017; Maurer et al., 2017). CA3 hyperactivity is suggested to bias the aged CA3 towards pattern completion, promoting maintenance of stored representations to result in rigidity of CA3 place fields (Wilson et al., 2006). However, age-related memory deficits are not only characterized by overly rigid representations. CA1 place cells in aged rats failed to stably retrieve the same map in a familiar environment (Barnes et al., 1997), suggesting that aged rats improperly remap in a familiar environment. Thus, the role of the hippocampus in the organization and representation of information may be compromised in aging, resulting in incorrect retrieval of the wrong maps for a given environment (Rapp, 1998; Redish et al., 1998; Tanila, 1998; Wilson et al., 2004).

Recent studies have shown compelling evidence for anatomical and functional heterogeneity along the CA3 transverse axis (Hunsaker et al., 2008; Ishizuka et al., 1990; Lee et al., 2015; Lu et al., 2015; Marrone et al., 2014; Nakamura et al., 2013; Sun et al., 2017; Witter, 2007). In comparison with each other, proximal CA3 receives greater input from the DG, whereas distal CA3 receives greater input from the entorhinal cortex and from the CA3 recurrent collateral systems. Functional gradients along the CA3 transverse axis suggest that proximal CA3, in conjunction with the dentate gyrus, supports pattern separation and distal CA3, with its extensive, recurrent collateral system, supports pattern completion (Lee et al., 2015; Lu et al., 2015). Proximal and distal CA3 send topographically organized projections to distal and proximal CA1, respectively, to form functionally distinct, parallel processing streams along the transverse axis in the hippocampal circuit (Amaral and Witter, 1989; Ishizuka et al., 1995; Lee et al., 2020). While age-related memory impairments have been linked to CA3 hyperactivity and enhanced pattern completion (i.e. rigid representations), how the age-related CA3 impairments map onto the heterogeneity along the CA3 transverse axis has not been determined. To investigate how aged CA3 hyperactivity may affect the parallel processing circuit within the hippocampus, we examined the CA3 place cell firing rates of young and aged rats. We show that the age-related CA3 firing rates are hyperactive in the proximal region but not in the distal region, where they may be hypoactive instead.

## Results

Multielectrode recordings were made from neurons along the dorsal CA3 transverse axis (Fig. 1A) from 4 Young (Y), 7 Aged-Unimpaired (AU), and 7 Aged-Impaired (AI) rats. Locations between 0-40% of the proximodistal length were defined as proximal CA3, 40-70% as intermediate CA3, and 70-100% as distal CA3 (Lu et al., 2015). The aged rats were pretested in the Morris water maze and categorized as learning impaired (AI) or learning unimpaired (AU) based on pre-determined learning index scores (Gallagher et al., 1993; Haberman et al., 2011; Branch et al., 2019) (Fig. 1B). All rats were then trained to run clockwise on a circular track containing salient local texture cues in a curtained room containing salient global landmark cues (Fig. 1C).

**Figure 1.**
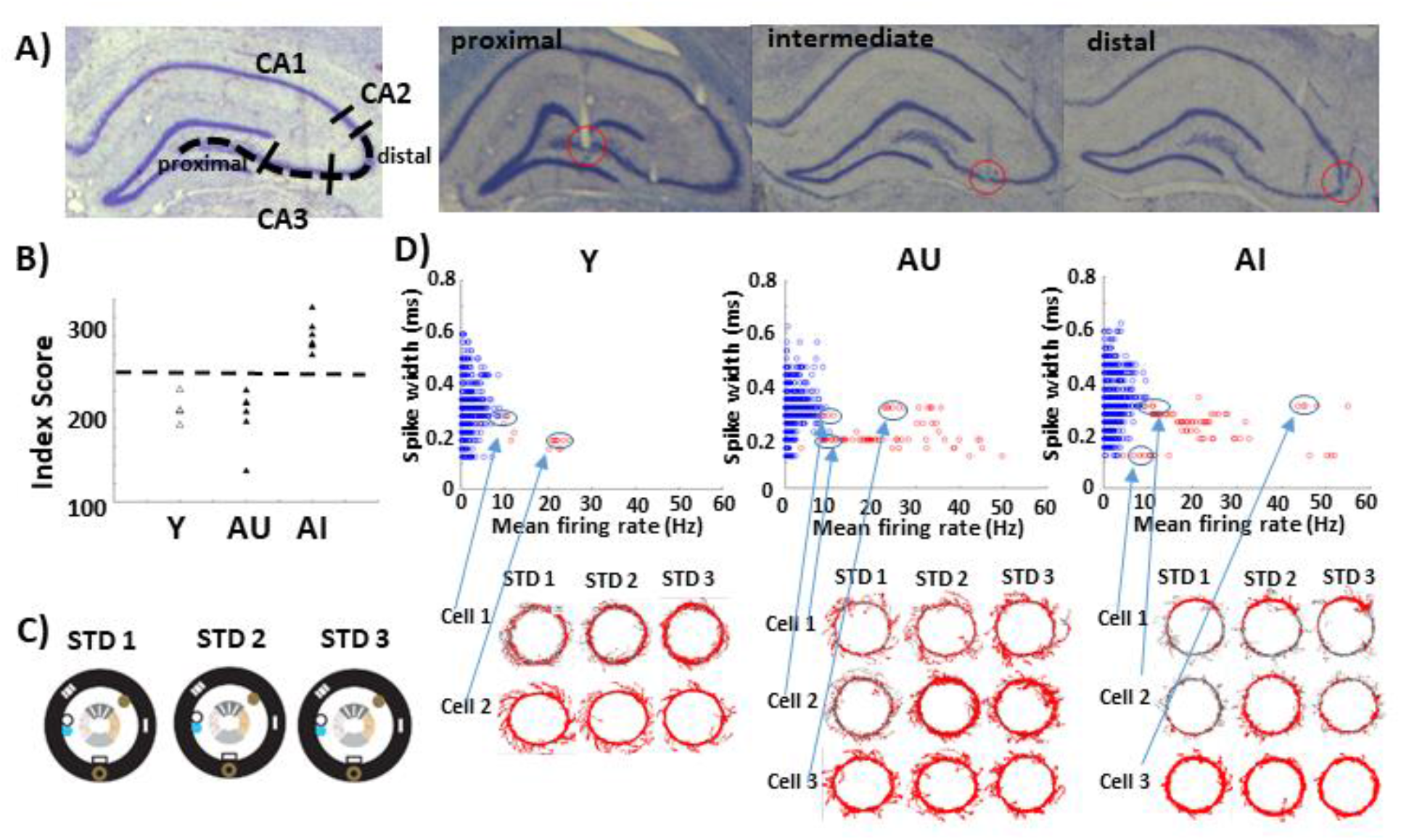
CA3 recording locations and cell classifications. **(A)** Recordings were made along the CA3 transverse axis. Locations between 0 and 40% of the axis length were defined as proximal, 40% and 70% as intermediate, and 70% and 100% as distal CA3 (left). Nissl-stained brain sections showing representative tetrode tracks in proximal, intermediate, and distal CA3 subregions (right). **(B)** A learning index score for each rat was derived during the probe trials during the water maze training, with lower scores indicating more accurate performance. Aged rats that performed on par with young rats were designated as aged-unimpaired (AU), and those that performed more poorly than young rats (index score >240) were designated as aged-impaired (AI). **(C)** Recordings were made during three standard (STD) sessions, in which the local and global cues were in a fixed configuration that the rat had experienced in all preceding training trials. **(D)** Classification of putative pyramidal cells and putative interneurons were made using three parameters: spike width, mean firing rate, and burst index. Using a K-means clustering analysis, blue circles represent cells classified as putative pyramidal cells and red circles represent cells classified as putative interneurons. Each circle is a cell recorded in one session. Hence, a cell can contribute up to 5 data points a day. Some of the classified putative interneurons had unusual firing patterns. Some cells in AU and AI rats had marked changes in their firing rates across sessions (AU cell 2; AI cell 2); one cell in the AI group had a very strong place field on the track (AI cell 1).

Using a K-means clustering analysis on three parameters (spike width, mean firing rate, burst index), cells were classified as putative pyramidal cells (low-rate, broad spikes, bursty) or putative interneurons (high-rate, narrow spikes, nonbursty) for each session (Csicsvari et al., 1999, 1998; Fox and Ranck, 1981, 1975). Figure 1D shows the classification for each cell-session (i.e., each cell can contribute up to 5 data points/day). There were few interneurons (red circles) recorded from Y rats, and they all showed the classic interneuron spatial firing pattern of being diffusely active at high rates along the entire track (Csicsvari et al., 1999, 1998; Fox and Ranck, 1981, 1975). In aged groups, some of the cells in the putative interneuron cluster did not seem like typical interneurons. Some fired at high rates but had broad spike waveforms more commonly associated with pyramidal cells (AU cell 3; AI cell 3). Such cells have been reported previously in CA1 from young adult rats (Csicsvari et al., 1999), and the lack of these cells in Y rats in the present data set likely is the result of the limited data set. However, some of the putative interneurons from aged rats had unusual spatial firing patterns. Whereas a typical interneuron will fire at similar rates in all sessions (but see (Wilent and Nitz, 2007)), some of the cells in AU and AI rats had marked changes in their firing rates across sessions (AU cell 2; AI cell 2). One cell in the AI group was classified as a putative interneuron because of its high firing rate and narrow spike, yet it had a very strong place field on the track (AI cell 1). Because of the limited sampling of these cells, no quantitative analyses have been performed and they are presented for descriptive purposes only. Thus, if a cell was classified as a putative interneuron cell (i.e., a high rate cell) in one of the sessions, that cell was classified as a putative interneuron for all five sessions and excluded from further analysis. Only well-isolated putative pyramidal cells (blue circles) were included in the analysis. Since the same cell can be counted up to 3 times (i.e., 3 STD sessions) in a given recording day, the values across all STD sessions in which a cell was active were averaged so that each cell contributed only one value to the analysis that day.

Most CA3 pyramidal cells are silent, or fire only rarely, in a given environment (Ahmed and Mehta, 2009; Alme et al., 2014), so we focused analysis on active pyramidal cells only. A well-isolated pyramidal cell was considered active if the cell fired > 50 spikes while the rat was running in a given session. An active cell was considered a place cell if it had a spatial information score > 0.75 bits/spike at a significance level p < 0.01 (Skaggs et al., 1996) and > 50 spikes when the rat’s head was on the track. As expected (Barnes et al., 1997, 1983; Robitsek et al., 2015; Shen et al., 1997; Tanila et al., 1997a, 1997b; Wilson et al., 2005), place cells were observed in all age groups across the 3 STD sessions (Fig. 2A). To examine place cell spatial stability, we analyzed a subset of place cells that were active in all three STD sessions. Place cells were ordered by the peak position of their linearized rate maps in the first STD session (STD1). Place field ordering remained stable in all three STD sessions in all age groups (Fig. 2B). To quantify the stability, we calculated the angular difference in the location of each place field on the circular track between STD1 and STD3. For all 3 age groups, the angular difference was centered around 0° with no significant age difference (Kruskal-Wallis test; *X*^2^[2] = 0.55, p = 0.76) (Fig. 2C).

**Figure 2.**
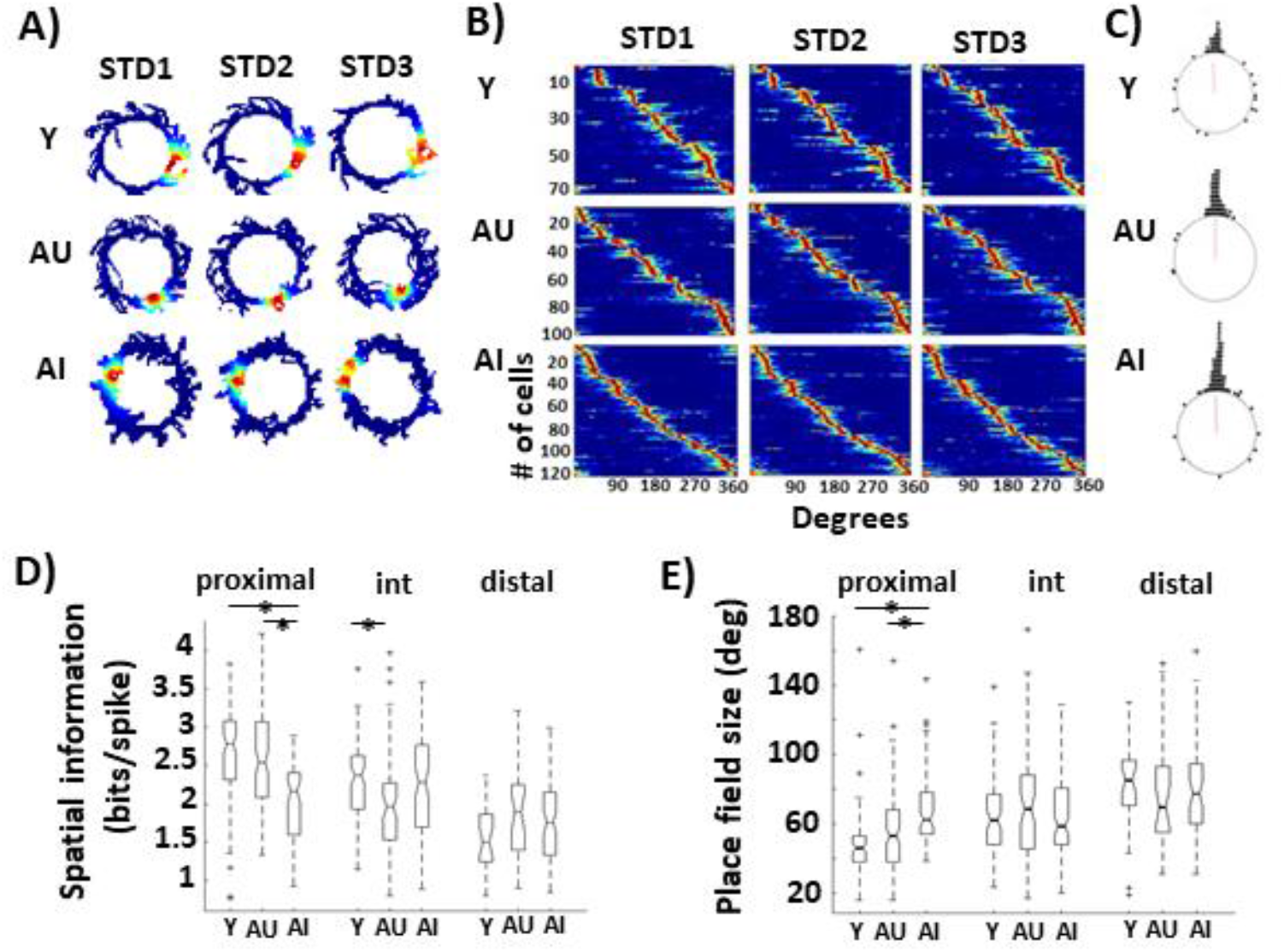
Place field properties in Y, AU, and AI rats. **(A)** Example cells recorded in all three STD sessions from Y, AU, AI rats. **(B)** From all four recording days, place cells that passed the inclusion criteria in all three STD sessions were ordered by the peak positions of their linearized rate maps in the first STD session (STD1). The positions of place fields that were active in all three STD sessions remained stable in all age groups. Firing rates of each place cell were normalized across the three sessions. The blue color indicates the minimum normalized firing rate (0), and the red color indicates the maximum firing rate (1). **(C)** The rotation amounts of the place fields between the first STD session (STD1) and the last STD session (STD3) were calculated. Comparisons of the rotation amounts did not show a significant difference across the age groups. **(D)** Spatial information scores and **(E)** place field sizes both showed significant group x region interaction effects. Both Y and AU rats showed decreasing spatial information scores from proximal to distal CA3 with corresponding increasing place field sizes from proximal to distal CA3. In AI rats, the spatial information scores and the place field sizes failed to show a normal gradient along the CA3 transverse axis. Post hoc Tukey’s tests (p < 0.05) showed a significant group difference in the proximal region, with AI rats showing lower spatial information scores and larger place field sizes compared to Y and AU rats.

Although place fields were stable in all groups, there were significant age differences in their spatial properties. Spatial information scores (Fig. 2D: Two-way ANOVA: group: F_(2,615)_ = 6.26, p = 0.002; region: F_(2,615)_ = 38.9, p < 0.0001; and interaction: F_(4,615)_ = 7.36, p < 0.0001) and place field sizes (Fig. 2E: Two-way ANOVA: group: F_(2,615)_ = 2.87, p = 0.057; region: F_(2,615)_ = 30.71, p < 0.0001; and interaction: F_(4,615)_ = 6.08, p < 0.0001) both showed significant group x region interaction effects. In Y and AU rats, the spatial information scores showed a decreasing trend from proximal to distal CA3, and correspondingly, the place field sizes show an increasing trend from proximal to distal CA3. These results are consistent with previous studies showing a decrease in spatial tuning from proximal to distal along the CA3 transverse axis in young rats (Lee et al., 2015; Lu et al., 2015). In contrast, the spatial information scores and the place field sizes of AI rats remained similar across the subregions. Post hoc Tukey’s tests (p < 0.05) showed a significant difference in the proximal region with AI rats showing lower spatial information scores with larger place field sizes compared to Y and AU rats, but no differences in distal CA3. These data suggest that AI rats do not show a normal gradient in spatial selectivity along the transverse axis, due to abnormally low spatial information in proximal CA3 but normal spatial selectivity in distal CA3.

### Aged CA3 place cells are hyperactive in proximal but possibly hypoactive in distal CA3

Young and aged CA3 place cells showed a gradient in mean firing rate along the CA3 transverse axis, but the gradients trended in opposite directions. While the young CA3 cells showed an increasing gradient in the mean firing rates from proximal to distal CA3 (as previously reported by Lee et al., 2015 and Lu et al., 2015), aged CA3 cells showed a decreasing gradient from proximal to distal CA3. Aged CA3 cells in the proximal region were hyperactive in both AI and AU rats but hypoactive in distal CA3 compared to Y rats (Fig. 3A: Two-way ANOVA: group: F_(2,615)_ = 0.83, p = 0.44; region: F_(2,615)_ = 1.96, p = 0.14; and interaction: F_(4,615)_ = 5.22 p = 0.0004; Tukey’s post hoc tests, p < 0.05).

**Figure 3.**
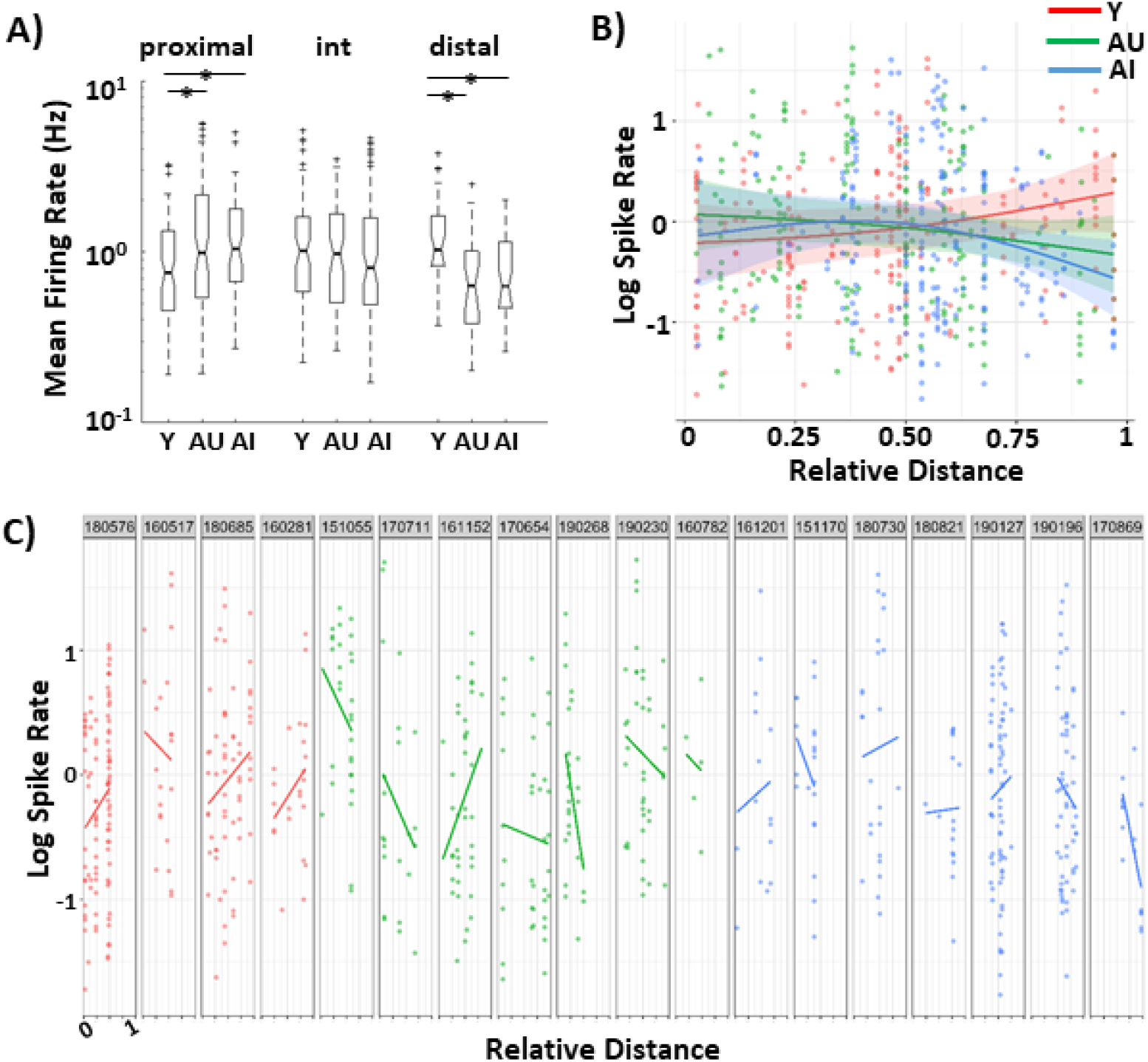
Age-related CA3 hyperactivity shows subregion differences along the transverse axis. **(A)** Mean firing rates of place cells showed a significant group x region interaction effect. Post hoc Tukey’s tests (p < 0.05) showed significant group differences in that aged CA3 cells were hyperactive in the proximal region but hypoactive in the distal region compared to young CA3 cells. **(B)** A linear mixed effects model was used to compare the mean firing rates along the transverse axis as a continuous variable, rather than a discrete subregion. Each circle (red = Y; green = AU; blue = AI) is a cell recorded at the location along the transverse axis. The solid lines show the model’s best fits, with 95% confidence intervals. Consistent with (A), both AU and AI show hyperactivity in the proximal region but hypoactivity in the distal region, compared with Y rats. **(C)** Data showing observed mean firing rates along the transverse axis organized according to each animal.

The data presented in Figure 3A were analyzed using standard methods from single-unit neurophysiology that typically treat each cell as an independent sample. However, certain assumptions of classic parametric statistical hypothesis-testing may be violated by these methods. (1) There may be correlations among the cells within an animal that violate the assumption of independence of each data point. (2) Data were taken across 4 days of recordings, and a typical level of instability of extracellular recordings makes it unknown which cells, and how many cells, were recorded more than once across the 4 days. Thus, the statistical power of these tests is artificially increased by an inflated sample size of unknown magnitude. (3) Inhomogeneities in sampling across animals (e.g., some animals may be biased toward proximal recording sites while others are biased toward distal sites) may also cause improper conclusions when the individual cells are pooled together. Because of these concerns, prior investigators have correctly argued that it is more conservative to use the individual animal as the unit of sampling for such statistical tests (Shen et al., 1997). However, such a conservative approach may reduce the power of the statistical tests, and it is often impractical to record from enough animals per group in these experiments, which require multiple months of effort per animal. Although preliminary analyses indicated that there was much greater variance among cells within an animal (93% of all variance) compared to variability across animals (7% of all variance), it was still possible that inclusion of a few outlier animals in one age group might skew the results. We thus used a linear mixed effects model (Laird and Ware, 1982) to account for the correlations among cells within an animal and the variability across animals to supplement the classic ANOVA tests. This analysis also allowed us to analyze location along the transverse axis as a continuous variable, rather than a somewhat artificial classification into discrete subregions (proximal, intermediate, distal).

Figure 3B shows the firing rates of each cell (log-scale ordinate, averaged across the 3 STD sessions of the day, as in Figure 2) for each tetrode location across all animals (abscissa). The lines show the model’s best fits. The average curves across animals differ significantly between the Y and AU groups (p = 0.03) and between the Y and AI groups (p = 0.01), but not between AU and AI groups (p = 0.61). Consistent with the ANOVA results, the AI and AU groups show a 19% hyperactivity in the more proximal region location (0.01 in Fig. 3B), the curves crossover midway through the transverse axis in the intermediate region, and the AI and AU groups show a 46% reduction in firing rate in the more distal regions (location 0.91). Figure 3C shows the data organized according to each animal, to show the variability in recording locations across subjects and the different relationships of firing rate to transverse axis location across animals that the linear mixed effects model takes into account.

### Speed does not account for the firing rate differences

Place cell firing rates are correlated with the animal’s running speeds (Czurkó et al., 1999; Ekstrom et al., 2001; Hirase et al., 1999; Maurer et al., 2005; McNaughton et al., 1983; Wiener et al., 1989). Because the aged rats ran at slower speeds on average compared to Y rats (Fig. 4A), speed can be a confounding variable that may contribute to the firing rate differences observed between the age groups. Thus, we ran another version of the same statistical model that incorporates momentary speed as a fixed effect that accounts for the potentially confounding effect of this variable. The results are similar to the model without the speed variable (Fig. 4B). The curves differ significantly between the Y and AU groups (p = 0.03) and between the Y and AI groups (p = 0.002), but not between AU and AI groups (p = 0.47). The hyperactivity of aged rats in proximal CA3 appears even stronger when speed is accounted for, compared to the previous model (Fig 3B), with the AI and AU groups showing a 72% hyperactivity in the more proximal region (location 0.01) and a 25% reduction in the more distal regions (location 0.91) compared to Y group. However, the hypoactivity of aged rats in distal CA3 appears weaker, and the crossover point is further toward the distal end. Based on the a priori definition of 0.7 being the transition between intermediate and distal CA3 (Lu et al., 2015), we tested whether the aged rats had statistically significantly lower firing rates than young rats from tetrodes located 70-100% along the transverse axis, but there was no statistically significant effect (95% CI: −5 to 41%; p = 0.14). We also performed a post hoc test on the data points 80-100% along the transverse axis, on the justification that prior work (Lu et al., 2015) showed a sharp transition, beginning around 80% of the transverse axis, in rate differences of cells in a global remapping experiment. From these data points, we estimated that Y rats had a firing rate that was 31% higher than the aged rats (95% CI: 4 to 64%; p = 0.02). Thus, although it is clear that proximal CA3 cells are hyperactive in aged rats and this hyperactivity diminishes in the more distal part of CA3, the results are suggestive, albeit somewhat ambiguous, as to whether the trend fully reverses to hypoactivity in distal CA3.

**Figure 4.**
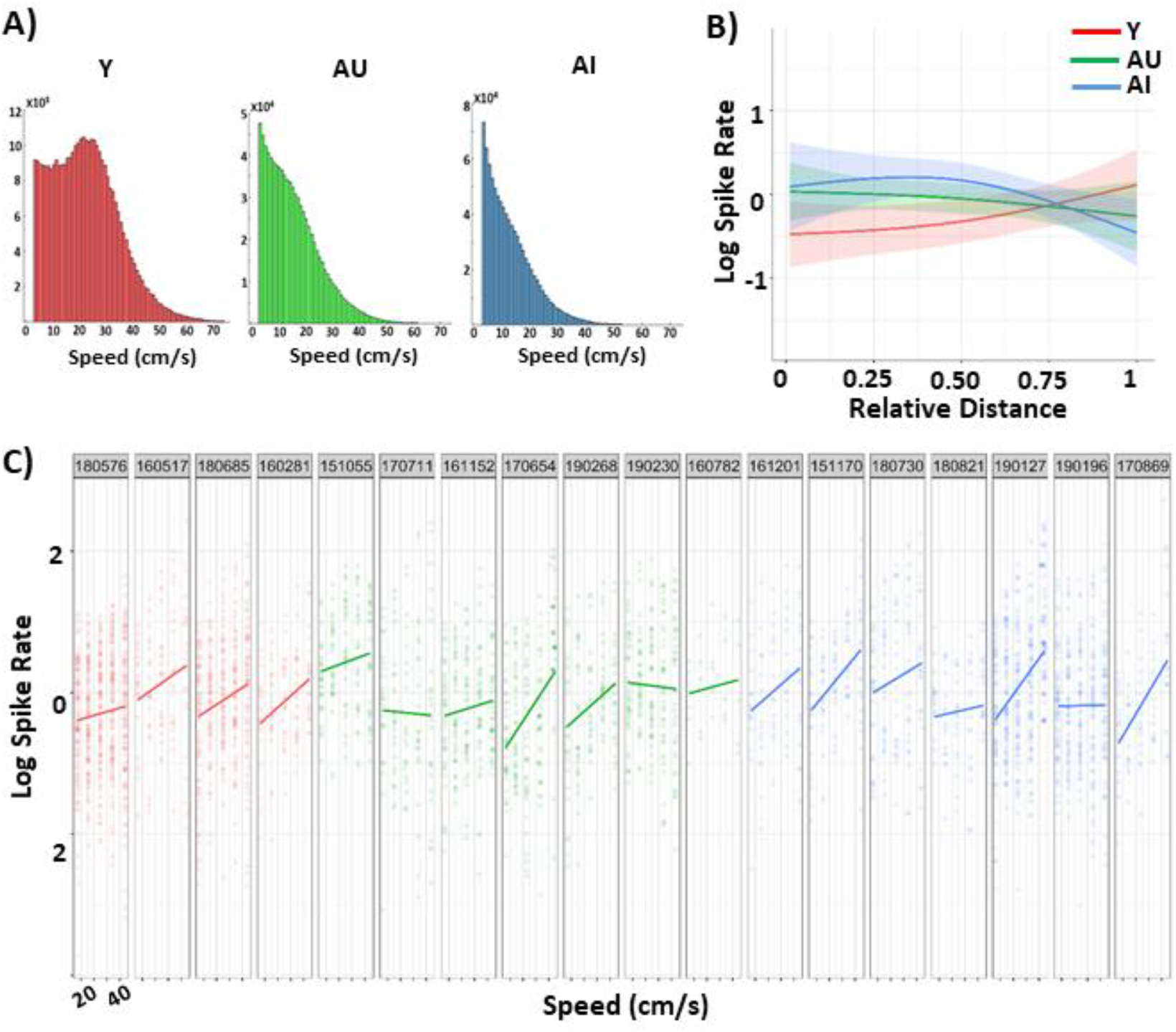
Speed does not account for the firing rate differences. **(A)** Histograms showing the distribution of speeds for each age group while animals performed the double rotation task. Aged rats ran at slower speeds compared to Y rats. **(B)** A linear mixed effects model that incorporates momentary speed as a fixed effect was used to analyze mean firing rate along the CA3 transverse axis. The solid lines show the model’s best fits, with 95% confidence intervals. With the confounding influence of speed taken into account, both AU and AI still showed hyperactivity in proximal CA3 but potential hypoactivity in distal CA3. **(C)** Data showing the observed mean firing rate relationship with the rat’s running speeds organized according to each animal.

Individual aged rats showed trends of more heterogeneous speed-firing rate relationships than young rats (Fig. 4C). Consistent with prior reports (Czurkó et al., 1999; Ekstrom et al., 2001; Hirase et al., 1999; Maurer et al., 2005; McNaughton et al., 1983; Wiener et al., 1989), all 4 Y rats showed increasing firing rates with increasing running speeds. However, the aged rats (especially AU rats) showed more heterogeneity in this pattern, as some aged rats showed the expected positive slopes whereas other rats showed negative or flat slopes.

### Fewer place cells are active in aged than young CA3

A large number of well-isolated, highly active pyramidal cells did not show localized firing locations on the circular track in aged rats (Fig. 5A). These active cells tended to fire either all over the circular track or during head-scanning behaviors, when the animals were looking out to the distal cues in the room (Monaco et al., 2014). To compare the proportions of nonspatial cells across the age groups, active cells were categorized into 3 groups: place cells (that meet spatial criteria: > 0.75 spatial information score with significance p < 0.01); low rate cells (that meet spatial criteria but have < 50 spikes when the rat’s head is on the track, i.e., excluding off-track spikes); and active, nonspatial cells (that do not meet the spatial criteria). There were significant group differences in the proportions of cell types in each subregion (Fig. 5B: proximal: Χ^2^= 10.87, p = 0.03; intermediate: Χ^2^ = 48.4, p < 0.0001; distal: Χ^2^ = 9.53, p = 0.049). AI rats had a lower proportion of place cells and a higher proportion of nonspatial cells in all the subregions compared to Y rats. The previous analyses were limited to cells that were classified as place cells, in accordance with prior literature on hyperactivity of CA3 place cells in hippocampus (Robitsek et al., 2015; Wilson et al., 2005). Because there was a higher proportion of active, nonspatial cells in AI rats than the other groups, we examined whether the nonspatial cells contribute to the age-related hyperactivity or hypoactivity. The average firing rates of the active, nonspatial cells across the age groups were not significantly different, except in the distal region where AU rats had lower rates compared to Y and AI rats (Fig. 5C: Two-way ANOVA: group: F_(2,279)_ = 0.82, p = 0.44; region: F_(2,279)_ = 3.44, p = 0.03; and interaction: F_(4,279)_ = 2.83 p = 0.03; Tukey’s post hoc tests, p < 0.05). The linear mixed effects model that incorporates momentary speed as a fixed effect on the active, nonspatial cells (as in Fig. 4B) showed no significant effects. Thus, these results suggest that although there are more active nonspatial cells in AI rats than the other groups, these cells do not contribute substantially to the age-related differences in CA3 firing rates; rather, only the place cells contribute to the age-related firing rate differences.

**Figure 5.**
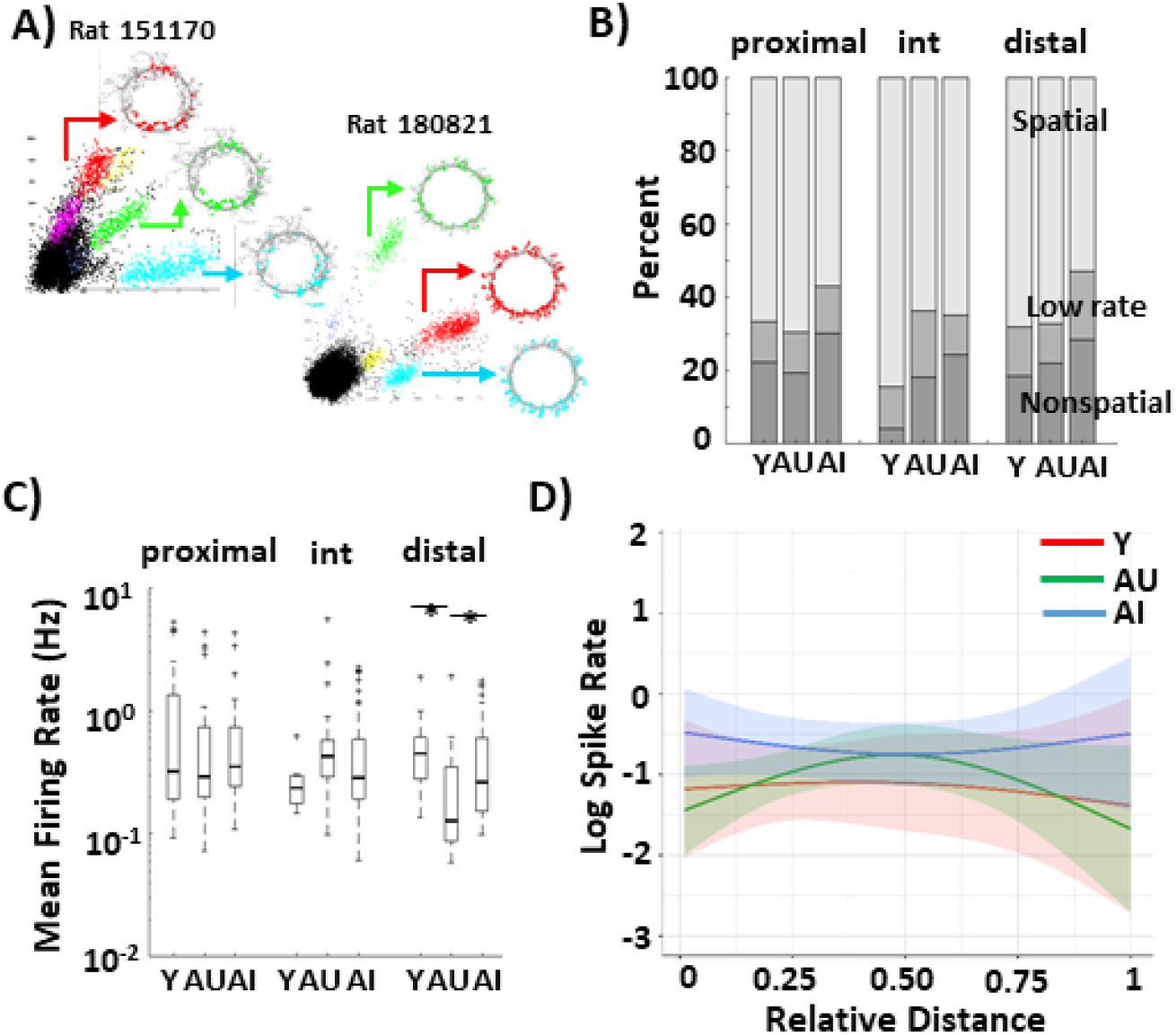
Nonspatial cells do not contribute to age-related hyperactivity. **(A)** Examples of well-isolated pyramidal neuron clusters from two different AI rats showing a lack of location specific firing on the circular track. **(B)** Active cells were categorized as spatial cells, low rate cells, or nonspatial cells. There were significant group differences in the proportions of cell types in each subregion (Χ^2^, p < 0.05). The proportion of active cells with spatial fields is lower in AI rats compared to AU and Y rats. Correspondingly, the proportion of nonspatial cells in AI rats was higher compared to AU and Y rats across the subregions. **(C)** Mean firing rates of nonspatial cells showed significant group x region interaction effects, but post hoc Tukey’s tests showed that the only difference was in the distal region where AU rats had lower rates compared to Y and AI rats. **(D)** A linear mixed effects model that incorporates momentary speed as a fixed effect was used to analyze the mean firing rates along the CA3 transverse axis. The solid lines show the model’s best fits, with 95% confidence intervals. The mean firing rates along the transverse axis were not significantly different among the groups.

## Discussion

The current study compared the firing properties of CA3 place cells from young and aged rats. CA3 place cells in aged rats (both AU and AI) were hyperactive in proximal CA3, which is consistent with the prior studies showing age-related hyperactivity in CA3 (Haberman et al., 2017a; Maurer et al., 2017; Robitsek et al., 2015; Wilson et al., 2005). Furthermore, our results showed some evidence that aged CA3 place cells were hypoactive, or at least different from young, in distal CA3. The firing rate differences we have shown along the CA3 transverse axis in aged rats may seem to contradict the Maurer et al. (2017) results, which showed that aged-poor performing rats had significantly higher numbers of *Arc*-positive cells expressing the immediate early gene *Arc* in CA3, but with no significant differences between proximal and distal CA3 subregions. However, the location of distal CA3 analyzed by Maurer et al. (2017) was not at the extreme distal end of CA3 but was located more toward the border of distal/intermediate CA3 (Figure 4 in Maurer et al., 2017). Furthermore, the two studies measured related, but fundamentally different, quantities. Although *Arc* studies are useful indicators of gross measurements of cell activity and the numbers of cells that are active under certain conditions (Guzowski et al., 1999), they are not accurate indicators of neural firing rates per se (Guzowski et al., 2006). The present study used single-unit recordings to measure the firing rates of CA3 place cells and showed the gradient in firing rate changes along the entire CA3 transverse axis, rather than just at a selected image region. Thus, there are no clear contradictions between the two studies. Instead, contrary to notions that there is an overall hyperactivity in aging CA3, our results demonstrate that age-related firing rate changes may be heterogeneous along the CA3 transverse axis, providing new insights to understanding the role of CA3 in age-related cognitive impairment.

### Role of aging CA3 in pattern separation and pattern completion

Classic computational models propose that the hippocampus performs two processes that maximize memory storage in a distributed network and minimize interference in memory retrieval: pattern separation, the ability to orthogonalize similar input patterns, and pattern completion, the ability to retrieve complete output patterns from partial or degraded input patterns. The dentate gyrus (DG) has been proposed to perform pattern separation (Marr, 1969; McNaughton and Nadel, 1990; Rolls and Treves, 1998; Yassa and Stark, 2011), while CA3, due to its recurrent collaterals, has been proposed to perform pattern completion (Marr, 1971; McClelland and Goddard, 1996; McNaughton and Morris, 1987; Rolls and Treves, 1998). However, there are anatomical and functional gradients along the CA3 transverse axis (Hunsaker et al., 2008; Ishizuka et al., 1990; Lee et al., 2015; Li et al., 1994; Lu et al., 2015; Marrone et al., 2014; Nakamura et al., 2013; Sun et al., 2017; Witter, 2007). Specifically, direct electrophysiological recordings of CA3 population activity demonstrated that proximal CA3 is biased toward pattern separation and distal CA3 is biased toward pattern completion (Lee et al., 2015; Lu et al., 2015).

Prior work suggested that the hippocampal system tradeoff between pattern separation and pattern completion might favor pattern completion in aged rats as the result of the hyperactivity in the pattern completion networks of CA3 of aged rats. This explanation fit with experimental data showing that AI rats were less likely than Y rats to form new representations when environments were altered (Robitsek et al., 2015; Tanila et al., 1997a, 1997b; Wilson et al., 2005, 2004). It also fit with data from humans showing that aged individuals had deficits in remembering whether an image of an object was a previously seen image (a target) or one that was slightly altered (a lure) (Bakker et al., 2015, 2012; Reagh et al., 2018). The authors interpreted these studies as a deficit in pattern separation, as the hyperactive CA3 networks tipped the balance in favor of pattern completion and prevented the memory networks from forming orthogonalized representations of the target and lure. Instead, the hyperactive CA3 caused the lure to recall the memory of the original target image. Treatment with the antiepileptic drug, levetiracetam, rescued the memory deficit, suggesting that tamping down the CA3 hyperactivity restored the balance between pattern separation and pattern completion (Bakker et al., 2015, 2012; Robitsek et al., 2015). Further work implicated hypoactivity in the LEC and the DG (Burke and Barnes, 2006; Geinisman et al., 1992; Reagh et al., 2018; Small et al., 2004) in the model. Because the LEC is especially disrupted in aging compared to the MEC (Reagh et al., 2018; Stranahan et al., 2011b, 2011a; Tran et al., 2019), and because the DG is thought to be crucial for performing normal pattern separation, the hypoactivity in these regions fit well with the notion that a major factor in age-related memory impairment is an imbalance between pattern separation and pattern completion. That is, a reduction in pattern separation caused by disruptions in the DG circuitry, coupled with an increase in pattern completion caused by increased CA3 activity, produces difficulties in subjects’ ability to create orthogonalized memory representations (especially for similar experiences), resulting in increased errors in memory recall.

Implicit in the former model is the notion that hyperactivity is present throughout the CA3 network. However, the present data are somewhat inconsistent with this view, because the part of CA3 that appears to underlie pattern completion (distal CA3) does not show hyperactivity in aged rats, and may even show hypoactivity. Instead, the part of CA3 that shows hyperactivity is the region that, along with the dentate gyrus, is involved in pattern separation (GoodSmith et al., 2019; Lee et al., 2020, 2015; Lu et al., 2015). Thus, using the logic of Wilson et al. (2006), a simple model might suggest that aged subjects should show greater pattern separation due to the hyperactivity specific to proximal CA3. However, considering the complex anatomy and parallel processing streams throughout the hippocampal transverse axis (Lee et al., 2020), we propose a revised model that presents a more nuanced view of how the patterns of hyperactivity and potential hypoactivity along the transverse axis of aged rats may result in an imbalance of the pattern-completion/pattern-separation balance. In this revised model, the bias toward pattern completion in aged rats results less from an enhancement of pattern completion, as suggested by Wilson et al. (2006), but rather from a weakening of pattern separation.

CA3 receives three major inputs (Fig. 6A): (1) perforant path inputs from layer II of the entorhinal cortex (Witter, 1993); (2) mossy fiber inputs from the DG (Blackstad et al., 1970; Swanson et al., 1978); and (3) recurrent collaterals from CA3 (Amaral and Lavenex, 2007). There is an increasing gradient of entorhinal inputs along the transverse axis from distal to proximal that is mirrored by a decreasing gradient of mossy fiber inputs (Sun et al., 2017). Thus, proximal CA3 receives the strongest DG inputs with minimal entorhinal inputs while distal CA3 receives the strongest entorhinal inputs (Claiborne et al., 1986; Ishizuka et al., 1990; Sun et al., 2017; Witter, 2007). In human and rat studies, aging is associated with neurofibrillary tangles (Braak and Braak, 1995), loss of cells in layer II by the time of a clinical diagnosis of Alzheimer’s Disease (Gallagher and Koh, 2011; Gómez-Isla et al., 1996), loss of synaptic markers, such as reelin (Stranahan et al., 2011a, 2011b), and hypoactivity (Reagh et al., 2018) in the entorhinal cortex. The entorhinal cortex is the major input to the DG. With reduced entorhinal function, the DG in aged rats receives fewer synaptic contacts from the entorhinal cortex than in young rats (Burke and Barnes, 2006; Geinisman et al., 1992), which may correspond with decreased activity in aged DG (Small et al., 2004). In contrast, disruptions of inhibitory circuits in the aged DG (Andrews-Zwilling et al., 2012, 2010; Spiegel et al., 2013; Tran et al., 2019) have the potential to increase the output of DG neurons, perhaps disrupting pattern separation by decreasing the sparsity of the DG population. Thus, because proximal CA3 in aged rats presumably receives a weaker drive from pattern-separated outputs from DG, the recurrent collaterals in proximal CA3, facilitated by the hyperactivity in the aged rats (perhaps due to a reduction of feedforward inhibition from EC and feedback inhibition from decreased hilar interneurons with aging; (Andrews-Zwilling et al., 2012, 2010; Spiegel et al., 2013), may override the DG inputs and create a larger basin of attraction that produces pattern completion (Fig 6B,C).

**Figure 6.**
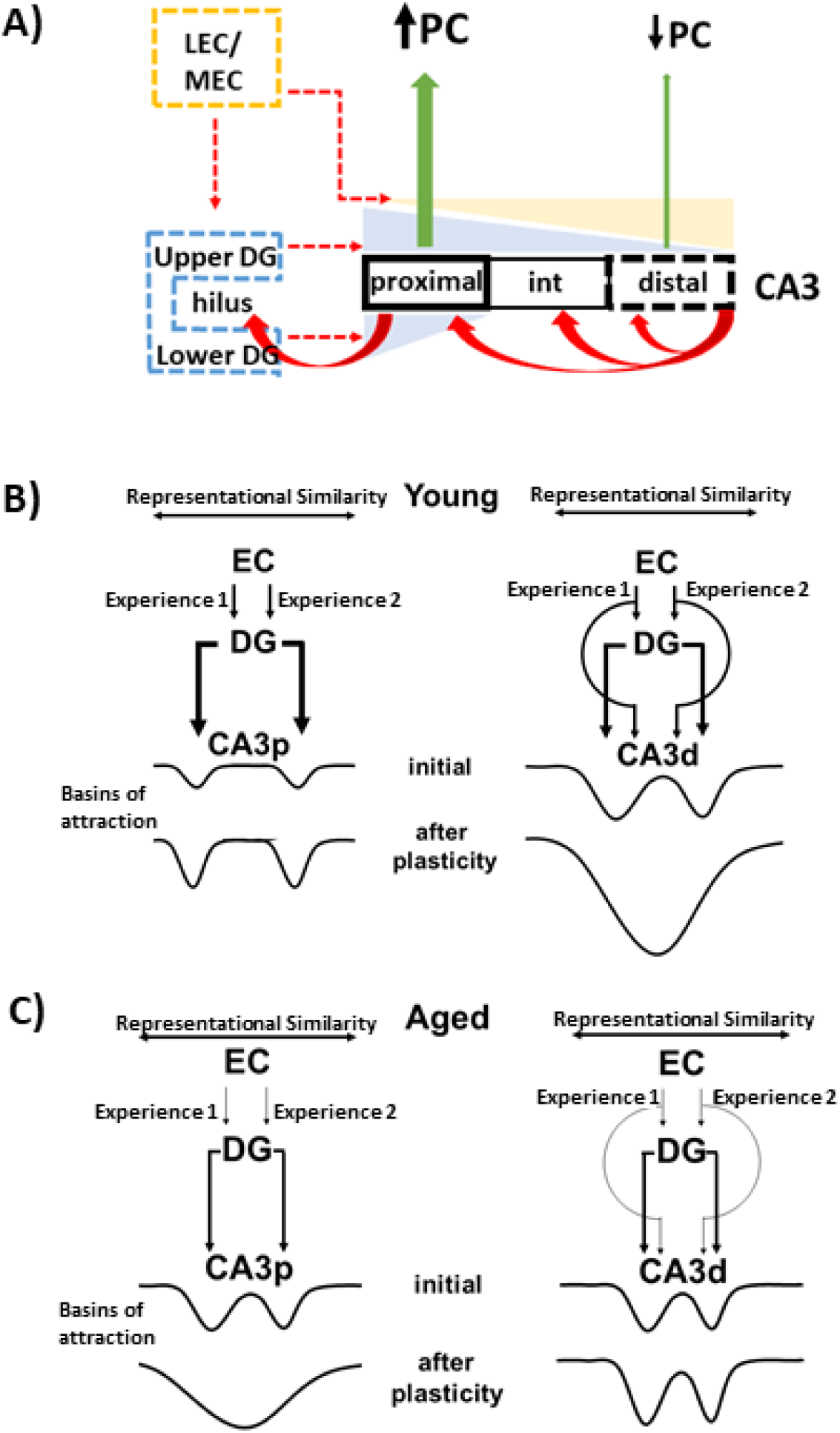
Hypothesized model of how changes in CA3 activity rates along the transverse axis affect the attractor dynamics of CA3 and the transmission of pattern-separated outputs from DG. **(A)** Three major inputs to CA3 are EC (with stronger inputs to distal CA3), DG (with stronger inputs to proximal CA3), and recurrent collaterals. Broken lines denote age-related reduction in function. In aging, with reduced EC and DG inputs, hyperactivity in proximal CA3 results in increased pattern completion (PC) while potential hypoactivity in distal CA3 results in weakened pattern completion. **(B)** For Young rats, the standard model of DG is that the EC neural representations of two similar experiences will be similar (shown as distance between two input arrows along a “representational similarity” axis). The DG takes the two similar EC patterns and performs a pattern separation function, increasing the distance between the representations of the two experiences (i.e., the distance between the arrows representing the DG output to CA3 is larger than the distance between the two EC inputs). In proximal CA3 (CA3p), the strong input from the DG (both upper and lower blades), coupled with the lack of strong, direct EC inputs, imposes the separated patterns on CA3 attractor networks. Through learning, the recurrent collaterals of CA3 increase the strength of the attractor basins. In distal CA3 (CA3d), the DG inputs are somewhat weaker and the direct EC inputs are stronger than CA3p. Thus, the combined DG and EC input may impose initial patterns on CA3d that are somewhat closer together and overlapping than those imposed in CA3p. The stronger recurrent collateral system of CA3d may override the external drive from DG/EC and merge the two attractor basins to form a broader and deeper energy well, resulting in pattern completion (or generalization) of the two input patterns. **(C)** For Aged rats, disruptions in the DG circuitry, due to dysfunction of the EC inputs (especially LEC) and the local inhibitory circuitry of the DG hilus, are hypothesized to reduce the ability of the DG to separate the similar EC input patterns (i.e., the distance between the DG inputs to CA3 is not as large as in Young animals, and the coordinated drive to CA3 may be reduced). The hyperactivity of CA3p cells may increase the relative strength of the CA3p attractor dynamics, overriding the DG inputs and causing a single basin of attraction (similar to that of CA3d in Young animals, but weaker because of the lower density of recurrent collaterals in CA3p). Thus, CA3p, which normally transmits a pattern-separated signal to distal CA1 in Young rats, instead transmits a pattern completed signal in Aged rats, as seen in some studies (Tanila et al., 1997a, 1997b). The possible hypoactivity of CA3d cells may result in weaker attractor basins in CA3d compared to Young rats, preventing the merging of the attractor basins that occurs in Young rats and in CA3p of Aged rats. Thus, two stable attractor states, with relatively lower energy barriers between them, may result in the bistable representations seen in other studies (Barnes et al., 1997).

In contrast to studies showing enhanced pattern completion in aged rats (Tanila et al., 1997a, 1997b), Barnes et al. (1997) showed that in familiar environments, young rats maintain the same map across repeated visits to the environment whereas aged rats maintain multiple maps of the same spatial context, switching between maps in an apparently arbitrary manner across visits. In other words, the Barnes et al. (1997) study suggests that aged rats pattern separate improperly while the Tanila et al. (1997a; 1997b) studies suggest that aged rats pattern complete improperly. Previous authors have discussed how these two sets of results can be reconciled (Rapp, 1998; Redish et al., 1998; Tanila, 1998). Here, we propose a modification of this reconciliation based on the gradients seen along the CA3 transverse axis. In short, failure of aged rats to form new maps in the Tanila et al. (1997a; 1997b) studies may be due to increased pattern completion (resulting from hypothesized dysfunction of the pattern separation mechanisms in DG) in proximal CA3, as described above (Fig. 6C). Moreover, the presence of multiple maps in aged rats in Barnes et al. (1997) study may be due to multiple, weaker attractor states (Redish et al., 1998) in distal CA3 related to the possible hypoactivity in the most distal part of CA3 (Rolls, 2019) (Fig. 6C). Although pattern completion and pattern separation are often presented as competing processes that produce a single output, we have argued previously that the gradients along the CA3 transverse axis might allow the system to output simultaneously a pattern-separated output (from proximal CA3 through distal CA1) and a pattern-completed output (from distal CA3 through proximal CA1) (Lee et al., 2020). Distal CA1 is associated with the LEC and proximal CA1 is associated with the MEC. Thus, memory deficits in aging may arise from a weakening of pattern separation (localized in the circuit from proximal CA3 through distal CA1 to deep LEC) as well as a weakening of attractor dynamics (localized in the circuit from distal CA3 through proximal CA1 to deep MEC). Of particular importance for humans, there appears to be a disproportionate increase in proximal CA3 compared to intermediate and distal CA3, as ~75% of CA3 appears to be homologous to rodent proximal CA3 (Lim et al., 1997). This disproportionate increase in proximal CA3 may explain why human fMRI studies have shown evidence for hyperactivity and stronger pattern completion in older adults’ DG/CA3 (Reagh et al., 2018; Yassa et al., 2011, 2010). The relative size of proximal CA3 in humans suggests that the evolution of human episodic memory required an expansion of the pattern-separating function of the DG-proximal CA3-distal CA1-deep LEC pathway. Perhaps not surprising, these are the components of the hippocampal system that are affected earlier and more profoundly in aging.

## Methods

### Subjects and Surgery

18 male Long-Evans rats (4 young; 14 aged) were used in this study. Male rats (retired breeders) were obtained at 9 months of age from Charles River Laboratories (Raleigh, NC) and housed in a vivarium at Johns Hopkins University until behavioral assessment in the water maze at 22-26 months of age. Young rats were obtained from the same source and were housed in the same vivarium and tested at 4-6 months of age. Prior to the onset of behavioral training, all rats were individually housed with ad libitum access to food and water during a 12-hour light/dark cycle, unless noted otherwise. All animal care, housing, and surgical procedures conformed to the National Institute of Health standards using protocols approved by the Institutional Animal Care and Use Committee at Johns Hopkins University.

All rats were prescreened for spatial learning ability in the Morris water maze as previously described (Branch et al., 2019; Gallagher et al., 1993) prior to the implantation of recording electrodes. Briefly, all rats were trained for 8 days (3 trials per day) in the water maze to locate a submerged escape platform that remained at the same location in the tank. Every sixth trial was a probe trial (no escape platform for the first 30 seconds), allowing assessment of search proximity. The learning index (LI), an average of weighted proximity scores obtained during probe trials (Gallagher et al., 1993), was used to classify animals as aged-impaired (AI) or aged-unimpaired (AU), with low scores reflecting a more accurate search. A learning index cutoff at 240 was used to segregate aged rats into unimpaired (LI < 240) and impaired (LI > 240). On day nine, all rats were given six trials to locate a visible platform above the water level to screen for nonspecific task impairment such as swimming behavior and escape motivation. Following water maze testing, all rats were placed on restricted food access to reduce their body weight down to 85% while they were given foraging sessions (20 mins per day for 10 days) in a cylindrical apparatus. These foraging sessions were given to accustom the rats to forage for chocolate pellets (Bio-Serv, Flemington, NJ) before the subsequent training on the double rotation track.

For hyperdrive implantation, a custom-built recording drive that contained 15 independently moveable tetrodes was surgically implanted over the right hemisphere. To optimize the drive placement, recordings were performed during the surgery to find the lateral edge of CA3, which served as a landmark for the mediolateral placement of the drive; the most lateral tetrode ranged from 3.7 to 4.2 mm lateral to bregma and 3.4 to 4.0 mm posterior to bregma. The tetrode array was configured as an oval bundle, approximately 1.2 mm in length and 0.9 mm in width, and encompassed a 3 x 6 array of tetrodes spaced approximately 300 μm apart. The array was implanted at an angle of ~35°, which is orthogonal to the longitudinal axis in the dorsal hippocampus and approximately parallel to the transverse axis of the hippocampus.

### Local-Global Cue Mismatch (Double Rotation) Manipulation

The double rotation environment consisted of a wooden circular track (76 cm outer diameter, 56 cm inner diameter) with salient local texture cues consisting brown sandpaper, pebbled gray rubber mat, tan carpet pad, and gray duct tape with white labeling tape stripes. The circular track was placed in a black curtained room with salient global cues consisting a cardboard circle with a white rectangular box on the floor below it, a blue surgical drape and white towel on an intravenous drip stand, a black and white striped card, a roll of brown wrapping paper, and a white card. Over the course of 2-3 weeks, the rats’ weights were reduced to 80-85% of free-feeding weights for the young rats and 75% for aged rats. All rats were trained to run clockwise on the circular track for chocolate pellets, which the local track cues and the global cues in a fixed configuration. The double rotation experiment began when the rats were able to consistently complete 5 training sessions per day with at least 10 laps/session. The double rotation experiments were conducted for 4 days, with 5 sessions each day. Three STD sessions (local and global cue relationships remained constant) were interleaved with two MIS sessions (local and global cues were rotated by equal increments but in opposite directions, producing mismatch angles of 45°, 90°, 135°, and 180°). Mismatch angles were chosen in pseudorandom order such that each angle was chosen once during the first 2 days of recording and once again during the second 2 days. On each recording day, baseline sessions (~15 min) in which the rat rested in the holding dish were recorded prior to the start and at the end of the experiment, in order to compare recording stability before and after the experiment. For this report, we analyzed cells only from the standard (STD) sessions to rule out any influence that may result from cue manipulations in the mismatch sessions.

### Electrophysiological Recordings

Tetrodes were made from 17 μm platinum-iridium wires (California Fine Wire Co., Grover Beach, CA). Impedance of the platinum wires was reduced to ~120 kOhms by electroplating them with platinum black. Neural signals were recorded using a 64-channel wireless transmitter system (Triangle Biosystems International, Durham, NC) and transmitted to a Cheetah Data Acquisition System (Neuralynx, Bozeman, MT). The signals were amplified 1,000-5,000 times and filtered between 600 Hz and 6 kHz (for units) or 1 and 600 Hz (for LFP). The spike waveforms above a threshold of 40-70 μV were sampled for 1 ms at 32 kHz, and LFPs were continuously sampled at 32 kHz. The rat’s position was tracked with an overhead camera recording light emitting diodes (LEDs) positioned over the head of the rat (red LEDs in front and green LEDs behind) at 30 Hz.

### Data Analysis

#### Unit Isolation

Multiple waveform characteristics (e.g., spike amplitude and energy) were used to isolate single units using custom-written, manual cluster-cutting software. Cells were assigned to subjective isolation quality scores from 1 (very good) to 5 (poor), depending on the distance each cluster was separated from the other clusters and from the background activity level. Cluster isolation was judged independent of the behavioral firing correlates of the cells. Three parameters were used to classify cell types: spike width, mean firing rate, and burst index. Using a K-means clustering analysis on these parameters, cells were classified as putative pyramidal cells (low-rate, broad spikes, bursty) or putative interneurons (high-rate, narrow spikes, nonbursty) for each session. If a cell was classified as a putative interneuron cell (i.e. a high rate cell) in one of the sessions, that cell was classified as an interneuron for all five sessions. Only well-isolated putative pyramidal cells (with isolation quality scores 1-3) were included in the analysis. Putative interneurons were excluded from the analysis.

Classified putative pyramidal cells were categorized as active cells if the cell fired more than 50 spikes while the rat was running (> 3 cm/s) on the track. Of the active cells, cells were categorized into 3 groups: place cells (meet spatial criteria: > 0.75 spatial information score with significance p < 0.01); low rate cells (meet spatial criteria but have < 50 spikes when the rat’s head is on the track); and active, nonspatial cells (do not meet the spatial criteria).

#### Rate Maps and Place Fields

The position and the head direction of the rat were based on tracking the LEDs on the headstage connected to the hyperdrive. Analysis was performed on data restricted to times when the animal’s head was within the boundaries of the circular track and to the animal was moving forward on the track at a speed > 3 cm/s. The x and y dimensions of the camera image (640 x 480 pixels) were divided into bin sizes of 10 pixels. A 2D firing rate map of a cell was generated by dividing the number of spikes of a single neuron in each bin by the amount of time the rat spent in that bin. For quantitative analysis, 2D rate maps were transformed into linear rate maps by converting the rat’s Cartesian position into units of degrees on the circular track; the linearized rate maps were used to calculate the spatial information score (Skaggs et al., 1996), the mean and the peak rates, and the place field size. Linearized rate maps were divided into 360 bins (1°/bin) and smoothed with a Gaussian filter (σ = 3°). Place cells were identified as neurons with spatial information scores > 0.75 bits/spike (Skaggs ref), spatial information significance p < 0.01, and number of on-track running spikes > 50. Spatial information significance was calculated with a shuffling procedure, in which the spike train and the position train were shifted relative to each other by a random time (minimum 30 s; the later time stamps at the end of the session were wrapped around to the beginning of the session), the rate map was recalculated, and a spatial information score was calculated. This procedure was performed 1,000 times, and the cell was considered significant at p < 0.01 if the true information score exceeded the values of more than 990 scores from the shuffling procedure. Place field boundaries were determined using a floor threshold set to 10% of the peak firing rate. Bins with firing rates greater than the floor threshold were tracked in each direction from the peak bin of the place field until four consecutive subthreshold bins were reached. The size of each place field was measured across those above-threshold bins. The peak firing rate was the maximum firing rate in the rate map, and the mean firing rate was calculated by dividing the number of running spikes by the duration of the session.

#### Rotational Analysis

The linearized rate map in the first standard session (STD1) session was correlated with the linearized rate map in the third standard session (STD3) session. The linearized rate map in the STD3 session was then circularly shifted in 1° increments and was correlated with the STD1 session rate map at each increment. The shift producing the maximum correlation was assigned as that place field’s rotation angle.

#### Histological Procedures

Rats were deeply anesthetized and perfused with 4% formaldehyde. Frozen coronal sections (40 μm) were cut and stained with cresyl violet. Images of the brain slices were acquired with an IC Capture DFK 41BU02 camera (The Imaging Source, Charlotte) attached to a Motic SMZ – 168 stereoscope. All tetrode tracks were identified, and the lowest point of the track was used to determine the recording location. The position of the tetrode along the CA3 transverse axis was normalized from 0 to 1, with the distal end as 1. Distance of the CA3 tetrode was measured manually from the proximal end and scaled by the total length of CA3.

#### Statistical Analysis

Statistical tests were calculated using Matlab (Mathworks, Natick, MA) and R (R Core Team (2013). R: A language and environment for statistical computing. R Foundation for Statistical Computing, Vienna, Austria. URL http://www.R-project.org/). For group comparisons, data were first log-transformed due to the highly skewed distributions of the raw data (although the original data are presented in figures). Two-Way ANOVAs with post hoc Tukey’s tests were then performed and considered significant at p < 0.05.

### Linear mixed effects models

The linear mixed effects regression models used to estimate, separately for the young (Y), adult unimpaired (AU) and adult impaired (AI) animals, the mean firing rate as a function of the neuron’s distance along the CA3 transverse axis with and without control for animal running speed. The response variable is the total spike count for a cell divided by its total number of frames. The key predictors are the age group of the animal and the location of the neuron along the transverse axis. To compare the mean rate of firing among the three groups of rats, a linear mixed effects model was used after log transforming the observed rates). The fixed effects in the model included two indicator variables for the three groups interacting with a smooth function of location (natural cubic spline with 2 degrees of freedom; (Hastie et al., 2009)). The random effects allow animals to have random intercepts and random linear slopes along the CA3 axis; the random effects account for autocorrelation among cells from the same animal so that the inferences about the group differences are valid.

We used a Wald test of the null hypothesis that the three groups shared the same firing rate curve as a function of location. The Wald test statistic was compared to a chi-square distribution with 4 degrees of freedom (number of groups −1 = 2) x (number of degrees of freedom for each curve= 2). We also calculated 95% confidence intervals for the predicted curves for each group that are linear contrasts of the regression coefficients. To control for confounding by running speed, each animal’s speed was then included in the fixed effects as a natural cubic spline with 3 degrees of freedom. Otherwise the two analyses are the same.

## Acknowledgments

We thank Robert McMahan, Andrew Sherwood, Nicholas Lukish, Arjuna Tillekeratne, and Kimberly Nnah for assistance in running the behavior experiments and histological procedures. Supported by grant P01 AG009973.

## Author Contributions

H.L: designed research; performed research; analyzed data; wrote the first draft; wrote the paper; edited the paper

Z.W: analyzed data; edited the paper

S.Z: designed analysis; edited the paper

M.G: edited the paper

J.J.K: designed research; designed analysis; wrote the paper; edited the paper

